# Antioxidant enriched fraction from *Pueraria tuberosa* alleviates ovariectomized-induced osteoporosis in rats, and inhibits growth of breast and ovarian cancer cell lines *in vitro*

**DOI:** 10.1101/2020.09.21.305953

**Authors:** Swaha Satpathy, Arjun Patra, Muhammad Delwar Hussain, Mohsin Kazi, Mohammed S Aldughaim, Bharti Ahirwar

## Abstract

*Pueraria tuberosa* (*P. tuberosa*), known as Indian Kudzu belongs to family Fabaceae and it is solicited as “Rasayana” drugs in Ayurveda. In the present study, we analyzed the efficacy an antioxidant enriched fraction (AEF) from the tuber extract of *P. tuberosa* against menopausal osteoporosis and breast and ovarian cancer cell lines. The AEF from *P. tuberosa* was identified by determining phenolic composition (total phenolic and flavonoid amount). Antioxidant property (*in vitro* assays) was also carried out followed by analysis of the AEF for its antiosteoporotic and anticancer potentials. Antiosteoporotic activity of AEF was investigated in ovariectomy-induced osteoporosis in rats and *in vitro* anticancer activity by MTT assay. Also, the GC/MS analysis of AEF was performed to determine various phytoconstituents. A docking analysis was performed to verify the interaction of bioactive molecules with estrogen receptors (ERs). Ethyl acetate fraction of the mother extract was proved as the AEF. AEF significantly improved various biomechanical and biochemical parameters in a dose dependent manner in the ovariectomized animals. AEF also controlled the increased body weight and decreased uterus weight following ovariectomy. Histopathology of femur revealed the restoration of typical bone structure and trabecular width in ovariectomized animals after AEF and raloxifene treatment. AEF also exhibited *in vitro* cytotoxicity in breast (MCF-7 and MDA-MB-231) and ovarian (SKOV-3) cancer cells. Further, genistein and daidzein exhibited a high affinity towards both estrogen receptors (α and β) in docking study revealing the probable mechanism of the antiosteoporotic activity. GC/MS analysis confirmed the presence of bioactive molecules such as stigmasterol, β-sitosterol, and stigmasta-3,5-dien-7-one. The observations of this study vindicate the potency of AEF from *P. tuberosa* in the treatment of menopausal osteoporosis and cancer.

## Introduction

World Health Organization defines osteoporosis as a decrease of bone mineral density (BMD) to greater than 2.5 standard deviations of the standard reference for BMD in young health women [1]. Osteoporosis deteriorates BMD, bone architectural structure and enhances the risk of fracture. In addition, osteoporosis causes severe problems to human’s quality of life, such as disability, loss of living ability, and even death [2]. Variation in bone forming (osteoblastic) and bone resorbing (osteoclastic) cell function causes osteoporosis [3]. Osteoporosis has the highest prevalence in senile people and severely affects about 50% of menopausal women worldwide. The expected adult population over 60 years in India by 2050 would be 315 million signifying more incidence of osteoporosis compared to 26 million in 2003 [4]. A decrease in the level estrogen is the key contributing feature for menopausal osteoporosis (MO) in women. The reduced estrogen causes diminished bone formation, enhanced bone resorption, and elevated production of proinflammatory cytokines such as IL-1, IL-6, IL-7, and TNF-α [5]. The occurrence of MO is increasing day by day because of deskbound life style, environmental vulnerability, amenorrhea, hormonal alterations, early inception of puberty and ovarian disorders [6, 7]. Furthermore, several studies have demonstrated oxidative stress as an imperative factor prevalence of MO as a shortage of estrogen declines the antioxidant defense, and this lowered antioxidant levels promote bone loss [8, 9]. Oxidative stress could reduce the life span of osteoblasts by inhibiting osteoblastic differentiation and promoting bone resorption by boosting development and activity of osteoclasts, thus causing osteoporosis. In MO, the activated osteoclasts produce reactive oxygen species like superoxides and rise in malondialdehyde level in blood. These oxidative stresses also contribute to bone loss in osteoporosis [4]. Antioxidants can be useful in the management of MO by normalizing the altered osteoblastic and osteoclastic functions [10].

Several drugs, such as estrogens, biphosphonates, and parathyroid hormone analogs are used for the inhibition and management of osteoporosis. They promote bone formation or decrease bone resorption or both [11]. However, these treatments comprise serious concerns related to their safety and efficacy. Estrogen therapy is not preferred in patients with hepatopathy and venous embolism. Also, the possibility of cancers (breast, cervical, ovary), heart disease, and stroke are high in long-term use of estrogen [12]. Long-term application of biphosphonates shows adverse effects such as osteonecrosis of the jaw and atypical femoral fractures [13]. Parathyroid hormone analogs are costly, with patients needing daily injection, and may cause adverse consequence like osteosarcoma [14]. Therefore, it is important to develop drugs from plant origin with that have a protective effect on bone loss with fewer side effects. These plant-derived estrogenic compounds are known as “Phytoestrogens” and are accepted worldwide as safe treatments [7]. The phytoestrogens mostly include isoflavones, isoflavanones, coumestans, flavanones, chalcones, and flavones [15]. A considerable number of plant drugs in the form of extracts, fractions, herbal preparations, and isolated molecules have been studied to prevent or control osteoporosis [16]. Although these plant derived remedies are helpful in the management of MO, they may produce the side effects of supplemental estrogen [17, 18]. Hence, a search for safe, cheap, and effective natural agents for the management of MO is required.

Different species of Pueraria such as *P. lobata, P. mirifica, P. candollei* var. mirifica have been studied as protective agents against bone loss [19–21]. *Pueraria tuberosa,* known as Indian Kudzu belongs to family fabaceae, is solicited as “Rasayana” drugs in Ayurveda. This plant is used in various Ayurvedic preparations, traditional management of a wide range of ailments, and explored scientifically for an array of pharmacological activities. The plant is a rich source of various secondary metabolites and contains phytoestrogenic compounds such as quercetin, genistein, and daidzein [22]. Despite the significant pharmacological and phytochemical potential, the antiosteoporotic activity of *P. tuberosa* has not been explored. Our objective was to identify an antioxidant enriched fraction (AEF) from the tubers of the plant, and to investigate the preventive effect of AEF in menopausal osteoporosis and anticancer activity.

## Materials and Methods

### Chemicals, reagents and kits

The following chemicals in high grade were obtained commercially or as a gift: 3-(4,5-dimethylthiazol-2-yl)-2,5-diphenyltetrazoliumbromide (MTT) (Sigma-Aldrich, St Louis, MO, USA); Raloxifene (Cipla Ltd., Goa, India); Phosphorous, calcium, alkaline phosphatase, tartrate-resistant acid phosphatase, total cholesterol, and triglyceride kits (Span Diagnostic Pvt. Ltd.); Dimethyl sulfoxide (DMSO) and phosphate buffer saline (PBS) (Mediatech Inc., Manassas, VA, USA); Xylazine (Indian Immunologicals Ltd., Hyderabad, India); Ketamine (Neon Laboratories Limited, Thane, India); Diclofenac (Troikaa Pharmaceuticals Ltd., Ahmedabad, India); Gentamicin (Abbott, Pitampur, India); DPPH (1,1-diphenyl-2-picrylhydrazyl) (HIMEDIA Co. Ltd., India) were procured.

### Extraction of plant material and fractionation

Tubers of *P. tuberosa* were collected from Bilaspur, Chhattisgarh, India, with the help of the traditional practitioners and authenticated through the ICAR-National Bureau of Plant Genetic Resources, Regional Station, Phagli, Shimla, India. A voucher specimen has been preserved in the Institute of Pharmacy, GGU, Bilaspur for future references. The fresh tubers were cut into small pieces and dried under shade, then coarsely powdered and stored in an air-tight container until further use. The coarse powder material was extracted with ethanol using soxhlet apparatus. The extract was concentrated under reduced pressure using a rotary vacuum evaporator. The concentrated extract was suspended in distilled water and successively fractionated by liquid-liquid partitioning with n-hexane, ethyl acetate and n-butanol. Finally, the remaining aqueous fraction was also prepared. All the fractions were dried and stored in air tight container until further use.

### Identification of antioxidant enriched fraction

The mother extract (ethanol extract, PT), n-hexane fraction (PT1), ethyl acetate fraction (PT2), n-butanol fraction (PT3) and aqueous fraction (PT4) were evaluated for the antioxidant potential (by DPPH assay, ABTS assay and finding total antioxidant capacity) and phenolic composition (by total phenolic and flavonoid content determination) to identify the best antioxidant enriched fraction (AEF).

#### DPPH assay

Scavenging of 1,1-diphenyl-2-picrylhydrazyl (DPPH) free radical of all the samples was measured spectrophotometrically [23]. Two milliliters of the samples of different concentrations were added to one milliliter of DPPH solution (methanolic, 0.2 mM). Methanol was used as a control in place of the samples. The solutions were kept at room temperature for one hour in the dark, and then the absorbance was measured at 517 nm. The potential of free radical scavenging was represented as the percentage inhibition of DPPH radical, and was calculated using the following formula. The concentration producing 50% inhibition (IC_50_) was also established.

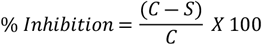

Where, C = absorbance of the control and S = absorbance of the sample

#### ABTS assay

Antioxidant capacity of the samples was analyzed based on their ability to interact with ABTS radicals [24]. The assay was performed following the protocol provided with the assay kit from Sigma-Aldrich, MO, USA (Catalog Number MAK187). The kit components were Cu^+2^ reagent (Catalog Number MAK187A), assay diluent (Catalog Number MAK187B), protein mask (Catalog Number MAK187C) and Trolox standard, 1.0 μmole (Catalog Number MAK187D). Briefly, 10 μL of the sample, 90 μL of HPLC water and 100 μL of Cu^+2^ working solution were transferred to each well in a 96 well plate. The contents were mixed thoroughly using a horizontal shaker and incubated in light protected condition at room temperature for 90 min. Finally, the absorbance was measured at 570 nm, and the Trolox equivalent as μM/g of the sample was determined from the standard curve of Trolox.

#### Determination of total antioxidant capacity (TAC)

TAC, in terms of copper reducing equivalent (CRE) of the sample, was evaluated using OxiSelect™ TAC Assay Kit (Cell Biolabs, Inc., San Diego, CA, USA; Catalog Number: STA-360) [25]. The components of the kit were uric acid standard (Part No. 236001), reaction buffer, 100X (Part No. 236002), copper ion reagent 100X (Part No. 236003) and stop solution, 10X (Part No. 236004). The assay protocol was as per the manufacturer’s product manual. Briefly, 20 μL of sample in various concentrations and 180 μL of 1X reaction buffer were transferred to each well in a 96 well plate and mixed thoroughly. An initial absorbance was taken at 490 nm. The reaction was started by adding 50 μL of 1X copper ion reagent into each well, and incubated on an orbital shaker for 5 min. Then the reaction was stopped by adding 50 μL of 1X stop solution to each well and absorbance was measured again. The net absorbance was calculated by subtracting the initial reading from the final reading and the mM uric acid equivalent (UAE) was determined from the uric acid standard curve. Finally the CRE was determined by multiplying UAE by 2189.

#### Determination of total phenolic and flavonoid content

Folin-Ciocalteu method and aluminum chloride colorimetric method were adopted for determining total phenolic content (TPC) and total flavonoid content (TFC), respectively [26] by reconstituting the samples in methanol. For determination of TPC, 100 μL of the sample (1.0 mg/mL) was mixed with 125 μL of Folin-Ciocalteu reagent and 750 μL of sodium carbonate solution (15% w/v) in a test tube. The final volume was adjusted to 5 mL with deionized water and mixed properly. The mixture was incubated at room temperature in the dark for 90 min, and then the absorbance was measured at 760 nm using a spectrophotometer. A blank sample with water and reagents was prepared and used as reference. TPC of the samples was represented as milligrams of gallic acid equivalents per gram dry weight (mg of GAE/g DW) of a sample through the calibration curve of gallic acid. For TFC estimation, 0.5 mL of sample (0.1 mg/mL) was mixed with 0.1 mL of AlCl_3_ (10%), 0.1 mL of potassium acetate (1 molar) and 1.5 mL of methanol (95%). The final volume was adjusted to 5 mL with distilled water and mixed thoroughly. The mixture was incubated in dark at room temperature for 60 min and then absorbance was measured at 415 nm. TFC was expressed as mg of rutin equivalents per gram (mg RE/g) of the sample through a standard curve of rutin. All measurements were carried out in triplicate.

### GCMS analysis of the AEF

Ethyl acetate fraction was identified as the antioxidant enriched fraction (AEF). GC/MS analysis was carried out on a GC/MS system comprising of Thermo Tracer 1300 GC and Thermo TSQ 8000 MS. The GC was connected to a MS with the following conditions such as TG 5MS (30m X 0.25 mm X 0.25 μm) column, operating in electron impact [electron ionization positive (EI+)] mode at 70 eV, helium (99.999%) as carrier gas at a constant flow of 1 ml/min, S/SL injector, an injection volume of 1.0 μl (split ratio of 10:1), injection temperature 250°C and MS transfer line temperature 280°C. The oven temperature was programmed from 60°C (isothermal for 2 min), with a gradual increase in steps of 10°C/min to 280°C. Mass spectra were taken at 70 eV, a scanning interval of 0.5 sec, and a full mass scan range from 50 m/z to 700 m/z. Data acquisition was carried out by Xcalibur 2.2 SP1 data acquisition software. Interpretation of the mass spectrum of GC/MS was performed by the NIST (National Institute Standard and Technology) mass spectral search program for the NIST/EPA/NIH mass spectral library version 2.0 g. NIST 11. The mass spectrum was compared with the spectrum of the components stored in the NIST library. The chemical name, molecular formula and molecular weight of the compounds were determined.

### Antiosteoporotic activity

#### Animals

Virgin female Wistar rats weighing 220-250 g were housed in polypropylene cages (two per cage) in air-conditioned room at 23±1 °C, relative humidity of 50-60% and 12 h/12 h light/dark illumination cycle. The animals were provided free access to diet and water. The experiment was performed after approval (Reference No.: 119/IAEC/Pharmacy/2015) by Institutional Animal Ethical Committee (IAEC) of Institute of Pharmacy, Guru Ghasidas University, Bilaspur, Chhattisgarh (Reg. No.: 994/GO/Ere/S/06/CPCSEA) under the guidelines of CPCSEA.

#### Acute oral toxicity study

An OECD 423 guideline was employed to determine the acute oral toxicity of AEF. The limit test was performed as per the guidelines on female rats (three rats per step) at a dose of 2000 mg/kg, orally and monitored for 14 days. The AEF was suspended in carboxy methyl cellulose (1.0%). Neither mortality nor any signs of moribund status were found at this dose (2000 mg/kg). Therefore, the LD_50_ cut-off is 5000 mg/kg (category 5 in the Globally Harmonized Classification System). The dosages selected for the antiosteoporotic property were 100 and 200 mg/kg/day.

#### Experimental protocol

The animals were acclimatized for seven days. On the seventh day, rats were ovariectomized and sham operated after anaesthetization with ketamine and xylazine intraperitoneally. The ovaries were bilaterally removed by a small midline skin incision and in the case of sham-operated group, the ovaries were exposed and sutured back without removing them [27]. Postoperative care was taken by administering diclofenac and gentamicin with individual housing of the animals for a few days. After four weeks, the animals were divided into different groups containing six animals each and treatment was continued for 90 days as below:

Group I: Sham-operated and received 1% CMC (Sham control).
Group II: Ovariectomized animals and received 1% CMC (OVX control).
Group III: Ovariectomized animals treated with standard drug, Raloxifene (1 mg/kg) (RAL).
Group IV: Ovariectomized animals treated with AEF (100 mg/kg) (AEF-100).
Group V: Ovariectomized animals treated with AEF (200 mg/kg) (AEF-200).

At the end of drug treatment, food was withheld for 24 h, and then the urine sample was collected in metabolic cages. Urine samples were refrigerated until further investigation.

Animals were sacrificed by ether anesthesia, and blood was withdrawn from the abdominal aorta. The blood samples were centrifuged at 2500 rpm for 25 min and stored for biochemical examination. Uterus was taken out watchfully after blood withdrawal and weighed. The femur and fourth lumbar vertebrae were collected by detaching the connecting tissue and stored at −70 °C until the biomechanical parameters were determined.

#### Determination of biochemical parameters

Various serum parameters were determined by using diagnostic kits. The parameters include calcium, phosphorus, alkaline phosphatase (ALP), tartrate resistant acid phosphatase (TRAP), triglycerides (TG), and total cholesterol (TC). Hydroxyproline (HP), calcium, and phosphorus in urine were also determined as reported earlier [4, 11].

#### Determination of biomechanical parameters

Weight (by digital balance), length (between the proximal tip of femur head and the distal tip of medial condyle) and thickness (using Vernier caliper) of femurs were measured after drying overnight and removal of bone marrow. Bone volume (by plethysmometer) and bone density (mass/volume) were also determined. The breaking strength of femur and fourth lumbar vertebrae was evaluated using hardness tester [11, 28].

#### Determination of body weight and organ weight

Bodyweight of each animal was measured on the first day and the last day of treatment. Uterus weight was also measured immediately after its removal and detachment of uterine horns, fat and connective tissues [11, 29].

#### Histopathology of femur

The right femur was fixed in 10% formalin for 12 h at 4°C, decalcified in ethylenediamine tetraacetic acid (EDTA) for 7 days, dehydrated, defatted, embedded in paraffin wax and section in the sagittal plane of 5 μm thickness was taken using a microtome. The sections were stained with hematoxylin and eosin (H & E), and scrutinized for histopathological changes under a light microscope (Primo Star, Zeiss with AxioCam ERc 5s camera) [7].

### *In vitro* cytotoxicity of antioxidant enriched fraction (AEF)

Breast (MCF-7 and MDA-MB-231) and ovarian (SKOV-3) cancer cells in DMEM media [supplemented with 10% fetal bovine serum (Mediatech, Manassas, VA) and 1% penicillin/streptomycin (Penicillin Streptomycin Solution 100X with 10,000 IU/mL penicillin and 10,000 μg/mL streptomycin, Mediatech, Manassas, VA)] were transferred to 96-well tissue culture plates at a density of 3000 cells/well, 24 h before treatment. The medium was then replaced with fresh medium containing AEF at various concentrations (31.5 to 500 μg/mL). The culture medium without any drug formulation was used as the control. After 72 h of incubation at 37°C and 5% CO_2_, media was removed carefully after taking out the culture plate from the incubator, and cells were washed twice with sterile PBS. 50 μl of MTT solution (0.5 mg/ml in DMEM media) was put into each well and further incubated for 4.0 h in the same condition. The medium was then removed from each well, and 100 μl of DMSO was added to each well to dissolve the purple formazan crystal obtained from the MTT assay. The absorbance value of each well was measured at 570 nm using a micro plate reader (Varioskan Flash, Thermo Scientific, USA). The percent cell viability with different treatments was calculated from the following formula [7, 30].

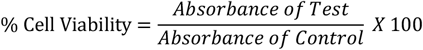

### Docking study of the identified phytoconstituents in AEF

In our earlier study, we have reported the presence of genistein and daidzein in the antioxidant enriched fraction (the ethyl acetate fraction) [31]. Docking study of these two compounds with estrogen receptor α (1×76) and estrogen receptor β (1×7R) [ER-α and ER-β] was performed to elucidate the mode of interaction. All computational studies were carried out using FlexX LeadIT 2.1.8 of BiosolveIT in a Machine running on a 2.4 GHz Intel Core i5-2430M processor with 4GB RAM and 500 GB Hard Disk with Windows 10 as the Operating System. The 3D conformer of the ligands was downloaded from PubChem in .sdf format. Reference protein coordinates of ER-α and ER-β for docking studies was obtained from X-ray structures deposited in Protein Data Bank (http://www.rcsb.org). For protein preparation, the chain having the receptor was selected as receptor components. Then reference ligand was selected. All the chemical ambiguities, which were crystallographically unresolved structures, were resolved, and the receptor was confirmed. The docking process deals with the translational, torsional, and ring conformation degrees of freedom. It was done by “Define Flex Docking” utility, and the FlexX accurately predicted the geometry of the protein-ligand complex within a few seconds. Then the docking was done using default parameters using a hybrid approach, followed by visualization using Pose View. The best conformation for each ligand sorted by the final binding affinity was stored [32].

### Statistical analysis

Data were represented as mean ± standard error means (SEMs). The data obtained in antiosteoporotic activity were subjected to a one-way analysis of variance (ANOVA) followed by post hoc Newman-Keuls multiple comparisons for significance using GraphPad Prism 7.0 (GraphPad Software, La Jolla, CA, USA) software. A value of p < 0.05 was considered as statistically significant.

## Results

### Characterization of antioxidant enriched fraction

The antioxidant potential of different samples (ethanol extract, and n-hexane, ethyl acetate, n-butanol and aqueous fractions) was evaluated based on their phenolic composition (total phenolic and flavonoid content as gallic acid equivalent and rutin equivalent, respectively), and antioxidant potential (by DPPH method, ABTS assay and determining total antioxidant capacity). Ethyl acetate fraction was regarded as the AEF for further study as it contained maximum phenolics and antioxidant activity (Tables 1 and 2).

**Table 1.** Total phenolic and flavonoid content, and antioxidant potential of ethanol extract and different fractions of *P. tuberosa*.

| Sample | TPC<br>(mg GAE/g DW) | TFC<br>(mg RE/g DW) | IC<sub>50</sub><br>(DPPH method)<br>(μg/mL) | μM Trolox equivalent/g<br>sample (ABTS assay) |
| --- | --- | --- | --- | --- |
| PT | 12.0 ± 0.70 | 40.26 ± 0.83 | 597.5 ± 7.89 | 376.13 ± 8.72 |
| PT1 | 0.95 ± 0.07 | 2.19 ± 0.35 | 1396.72 ± 15.85 | 45.87 ± 3.79 |
| PT2 | 106.23 ± 1.66 | 261.9 ± 1.73 | 55.70 ± 3.15 | 907.51 ± 8.07 |
| PT3 | 56.0 ± 1.30 | 25.56 ± 0.65 | 110.27 ± 10.41 | 360.26 ± 8.35 |
| PT4 | 26.46 ± 1.66 | 12.67 ± 0.77 | 291.08 ± 6.33 | 92.10 ± 4.84 |

Values are mean ± SEMs (n=3). PT, ethanol extract of *P. tuberosa*; PT1, n-hexane fraction; PT2, ethyl acetate fraction; PT3, n-butanol fraction; PT4, aqueous fraction; TPC, total phenolic content; TFC, total flavonoid content; GAE, gallic acid equivalent; DW, dry weight; RE, rutin equivalent; IC<sub>50</sub>, the concentration that provides a reduction of 50%; DPPH, 1,1-diphenyl-2-picrylhydrazyl; ABTS, 2,2'-azino-bis(3-ethylbenzothiazoline-6-sulfonic acid).

**Table 2.** Copper reducing equivalent of ethanol extract and different fractions of *P. tuberosa* at different concentrations.

| Concentration<br>(µg/mL) | Copper reducing equivalent |  |  |  |  |
| --- | --- | --- | --- | --- | --- |
|  | PT | PT1 | PT2 | PT3 | PT4 |
| 12.5 | 4.38±0.10 | 2.19±0.04 | 6.57±0.30 | 6.79±0.62 | 5.47±0.08 |
| 25 | 5.47±0.18 | 4.38±0.07 | 21.89±0.62 | 9.41±0.30 | 5.91±0.04 |
| 50 | 6.57±0.22 | 5.47±0.11 | 35.02±0.48 | 9.85±0.28 | 6.57±0.12 |
| 100 | 8.76±0.12 | 9.85±0.22 | 72.24±2.12 | 10.07±0.80 | 9.85±0.34 |
| 200 | 10.95±0.28 | 15.32±0.56 | 113.83±1.62 | 26.27±0.74 | 13.13±0.82 |

Values are mean ± SEMs (n=3); PT, ethanol extract; PT1, n-hexane fraction; PT2, ethyl acetate fraction; PT3, n-butanol fraction; PT4, aqueous fraction.

### GC/MS analysis of AEF

AEF from *P. tuberosa* contained 23 different chemical moieties (S1 as supplementary material) including stigmasterol, β-sitosterol and stigmasta-3,5-dien-7-one.

### Antiosteoporotic activity

The antiosteoporotic potential of AEF of *P. tuberosa* was evaluated in ovariectomized-induced osteoporosis in female rats by determining the following parameters:

#### Effect of AEF on biochemical parameters

Phosphorous (P) and calcium (Ca) level were analyzed both in serum and urine (Fig 1 and 2). OVX, as well as all other treatments did not significantly alter serum P and Ca. The level of P and Ca in urine increased significantly in the OVX group over sham control. Administration with both doses of AEF and raloxifene significant reduced the OVX-induced increase in urine P and Ca. Levels of bone markers, ALP and TRAP were significantly enhanced (p<0.001) after OVX.

**Fig 1.**
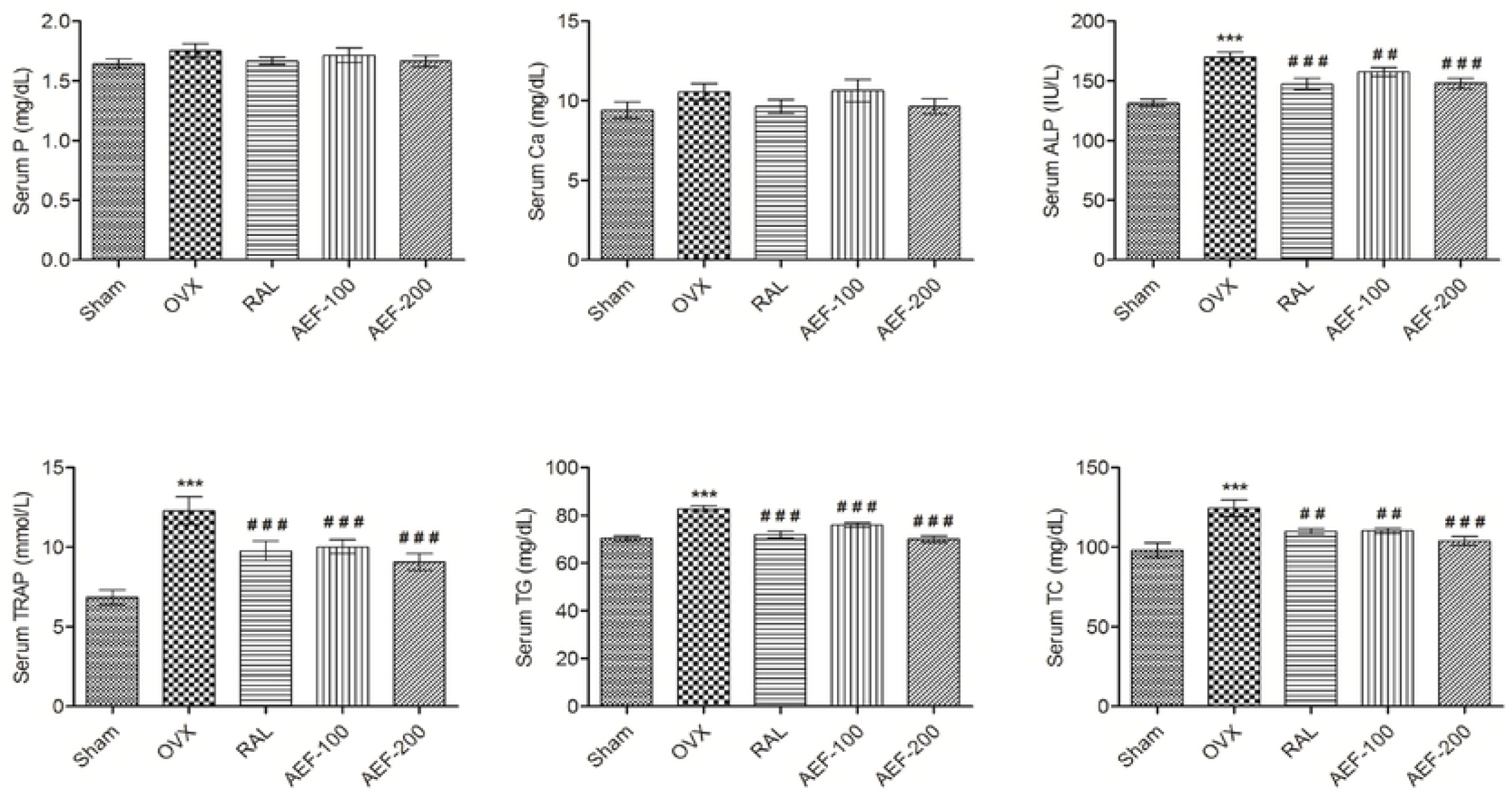
Effect of AEF from *P. tuberosa* on biochemical parameters of serum. Data were average ± SEM (n=6). *** p < 0.001 significantly different from sham control group. ## p < 0.01, ### p < 0.001 significantly different from OVX group. Ca, calcium; P, phosphorus; ALP, alkaline phosphatase; TRAP, tartrate resistant acid phosphatase; TG, triglycerides; TC, total cholesterol.

**Fig 2.**
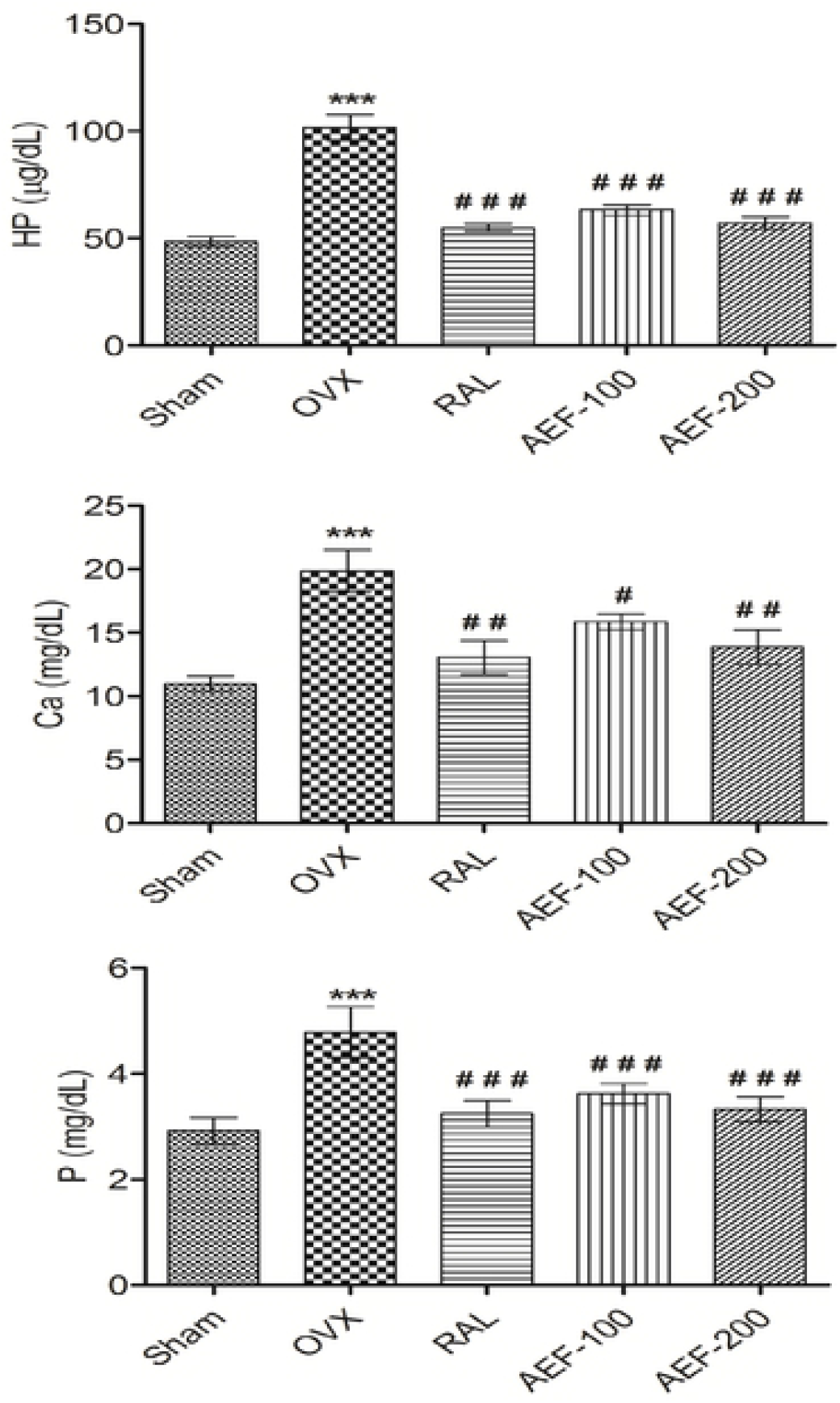
Effect of AEF from *P. tuberosa* on biochemical parameters of urine. Data were average ± SEM (n=6). *** p < 0.01 significantly different from sham control group. # p < 0.05, ## p < 0.01, ### p < 0.001 significantly different from OVX group. Ca, calcium; P, phosphorus; HP, hydroxyproline.

Both ALP and TRAP levels were reduced significantly and dose dependent after AEF treatment (versus OVX). Serum ALP and TRAP level were also reduced significantly after raloxifene treatment (Fig 1). OVX caused significant increase in the level of urine hydroxyproline (HP) compared to sham control. However the level of HP in raloxifene and AEF (100 and 200 mg/kg) treated groups was distinctly lowered (p<0.001) compared to the OVX group (Fig 2). Level of TC and TG increased significantly (p<0.001) in the OVX group compared to the sham control group (Fig 1). These increased TC and TG level was markedly lowered by AEF and raloxifene treatment. TG levels in raloxifene and AEF-200 groups are comparable with the sham control, and AEF exhibited better effect over raloxifene.

#### Effect of AEF on biomechanical parameters

None of the groups showed any significant alteration of femur length. Femur thickness, volume, weight and breaking strength were significantly decreased (p<0.001) in the OVX control compared to the sham control group. Significant increase in all these parameters (Fig 3 and 4) was observed with AEF and raloxifene administration. Furthermore, OVX caused a significant reduction of femur density, and treatment with AEF showed a substantial improvement of femur density. Treatment with raloxifene and AEF restored the breaking strength of 4^th^ lumbar vertebrae caused by ovariectomy (Fig 4).

**Fig 3.**
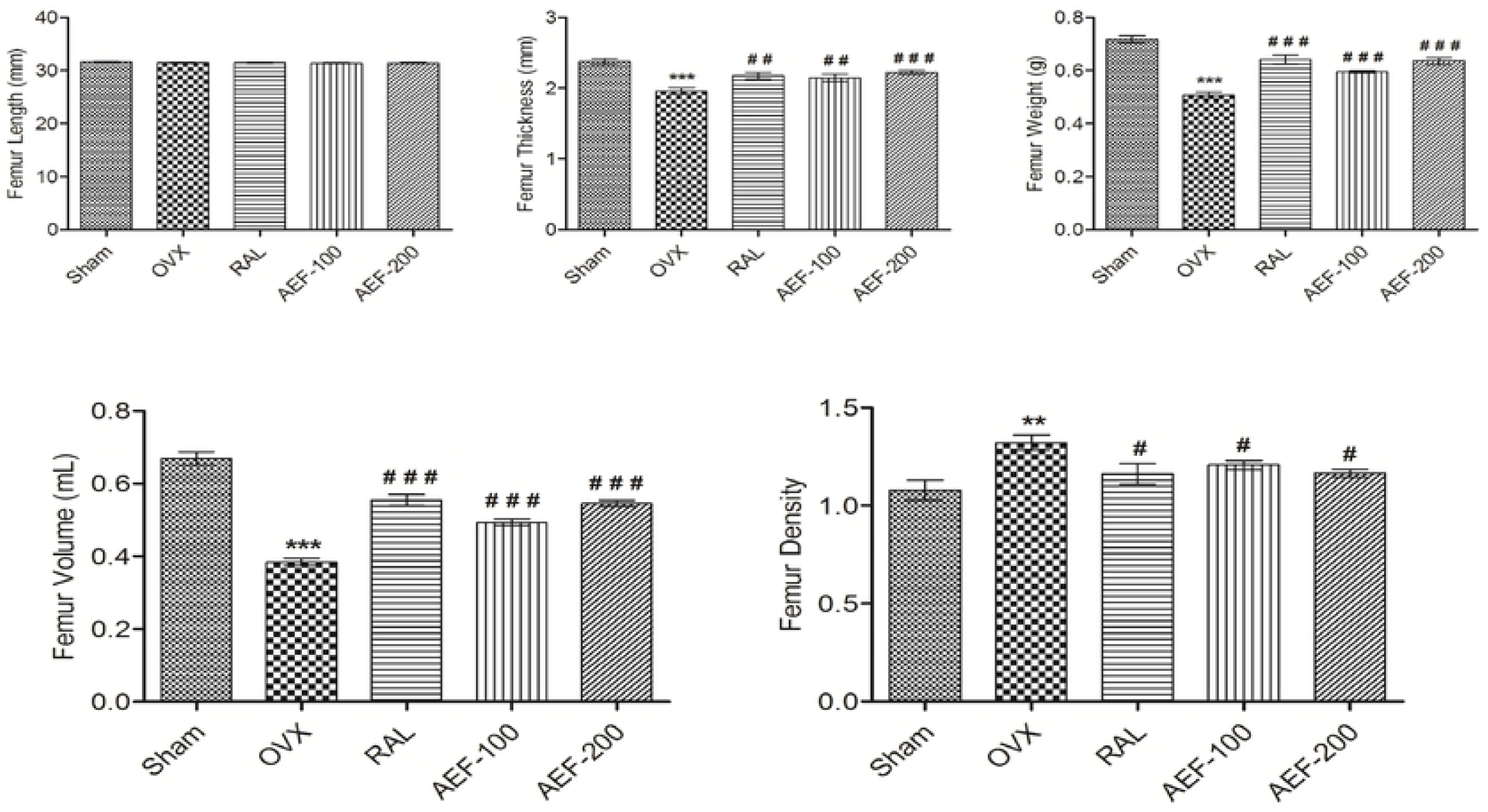
Effect of AEF from *P. tuberosa* on femur biomechanical parameters. Data were average ± SEM (n=6). ** p < 0.01, *** p <0.001 significantly different from sham control group. # p < 0.05, ## p < 0.01, ### p < 0.001 significantly different from OVX group.

**Fig 4.**
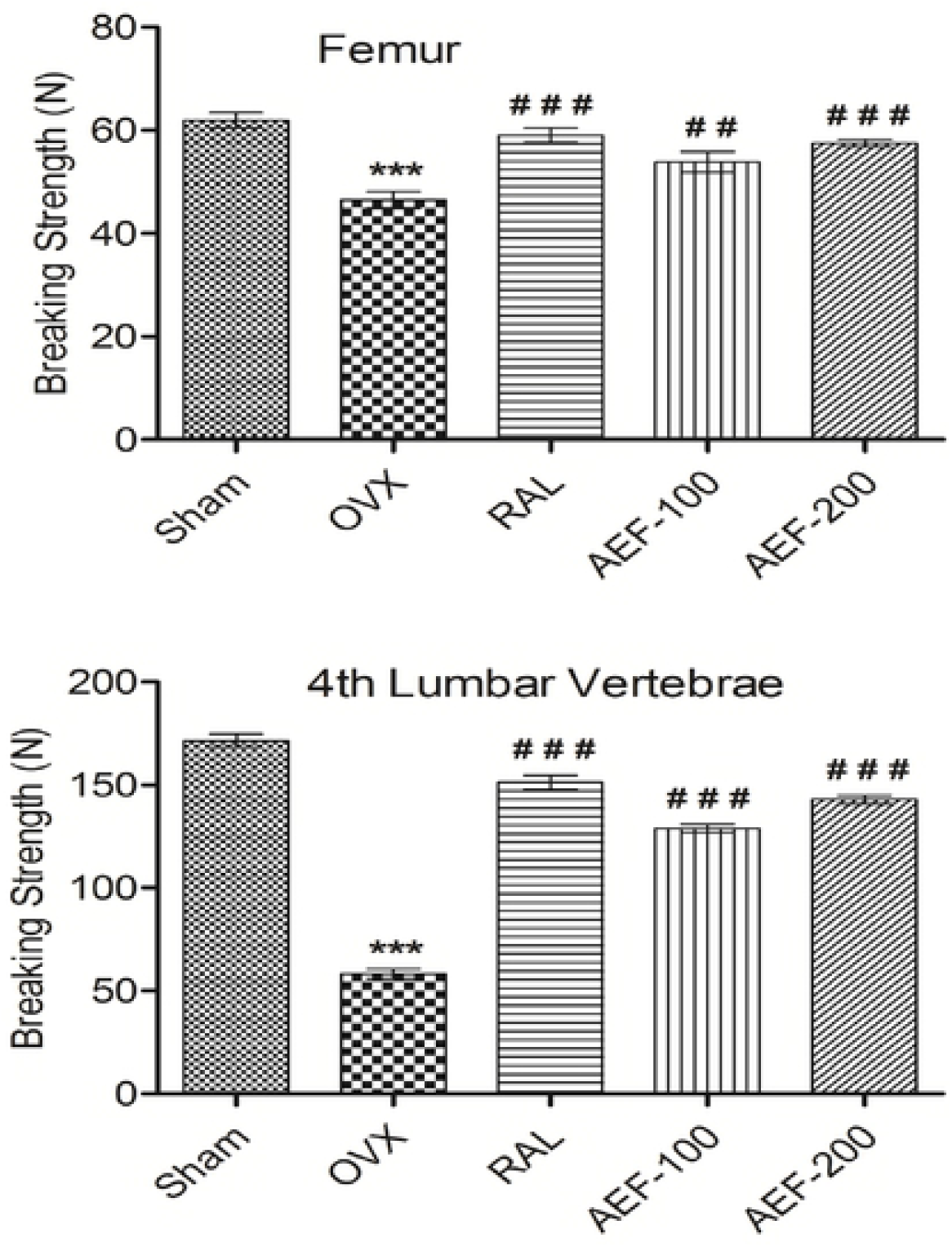
Effect of AEF from *P. tuberosa* on breaking strength of femur and 4^th^ lumbar vertebrae. Data were average ± SEM (n=6). *** p < 0.001 significantly different from sham control group. ## p < 0.01, ### p < 0.001 significantly different from OVX group.

#### Effect of AEF on body and organ weight

A significant (p<0.001) increase in body weight (BW) was observed due to OVX though there was no variation at the start of the study. Treatment with AEF and raloxifene markedly reduced the increased BW (Fig 5) as well as the final and initial BW difference compared to OVX. OVX caused a marked reduction in uterus weight. In comparison, administration of raloxifene, and AEF significantly increased uterine weight compared to OVX (Fig 5).

**Fig 5.**
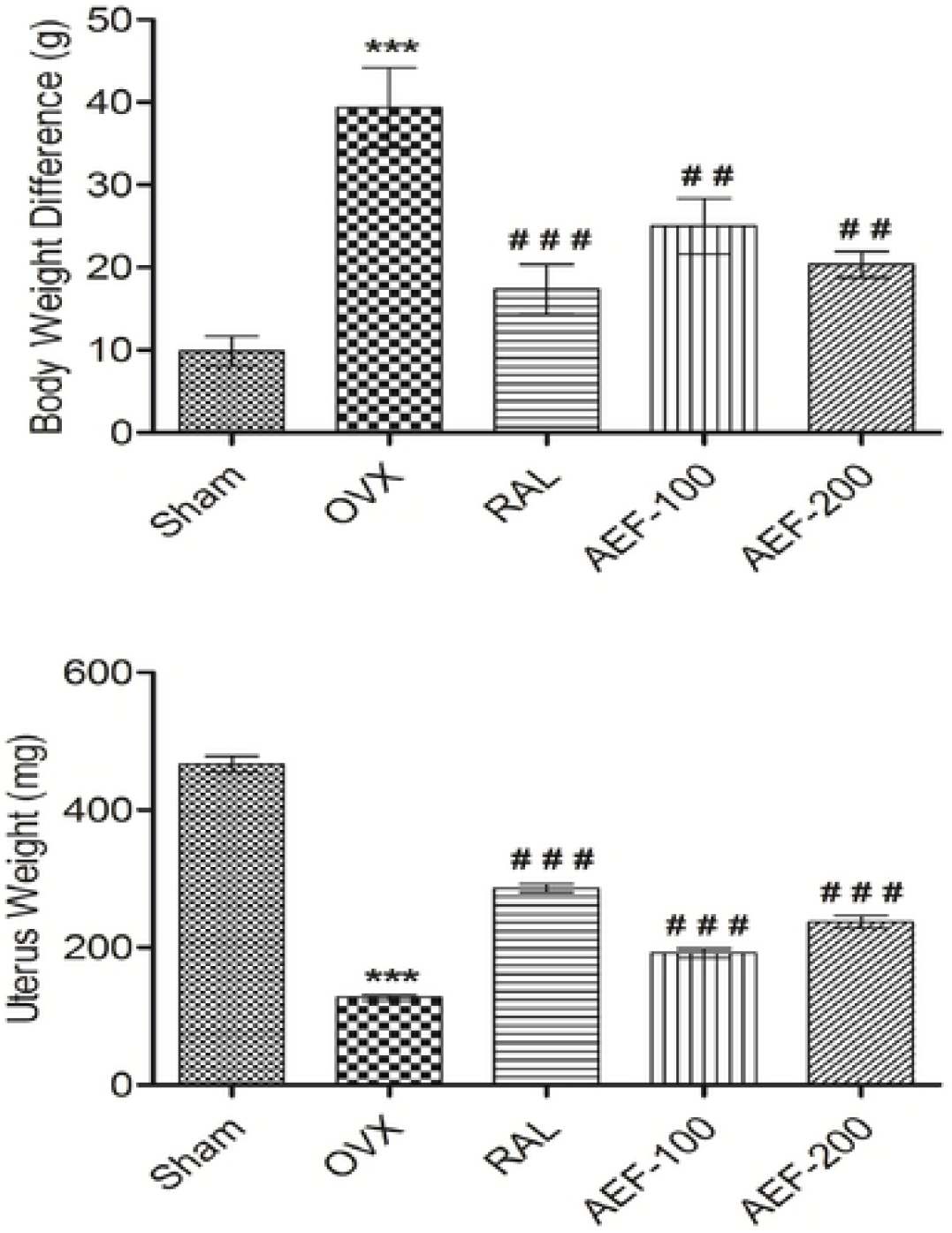
Effect of AEF from *P. tuberosa* on body and uterus weight. Data were average ± SEM (n=6). *** p < 0.001 significantly different from Sham control group. ## p < 0.01, ### p < 0.001 significantly different from OVX group.

#### Histopathology study

Photographs of the femur of different groups of animals are depicted in Fig 6A-E. There was a distraction of trabeculae with the decline in thickness and development of large cyst like spaces following OVX. Treatment with raloxifene and AEF showed trabecular ossification, mineralization, and compactness. Photomicrographs of raloxifene and AEF treated groups are indicative of the antiosteoporotic activity.

**Fig 6.**
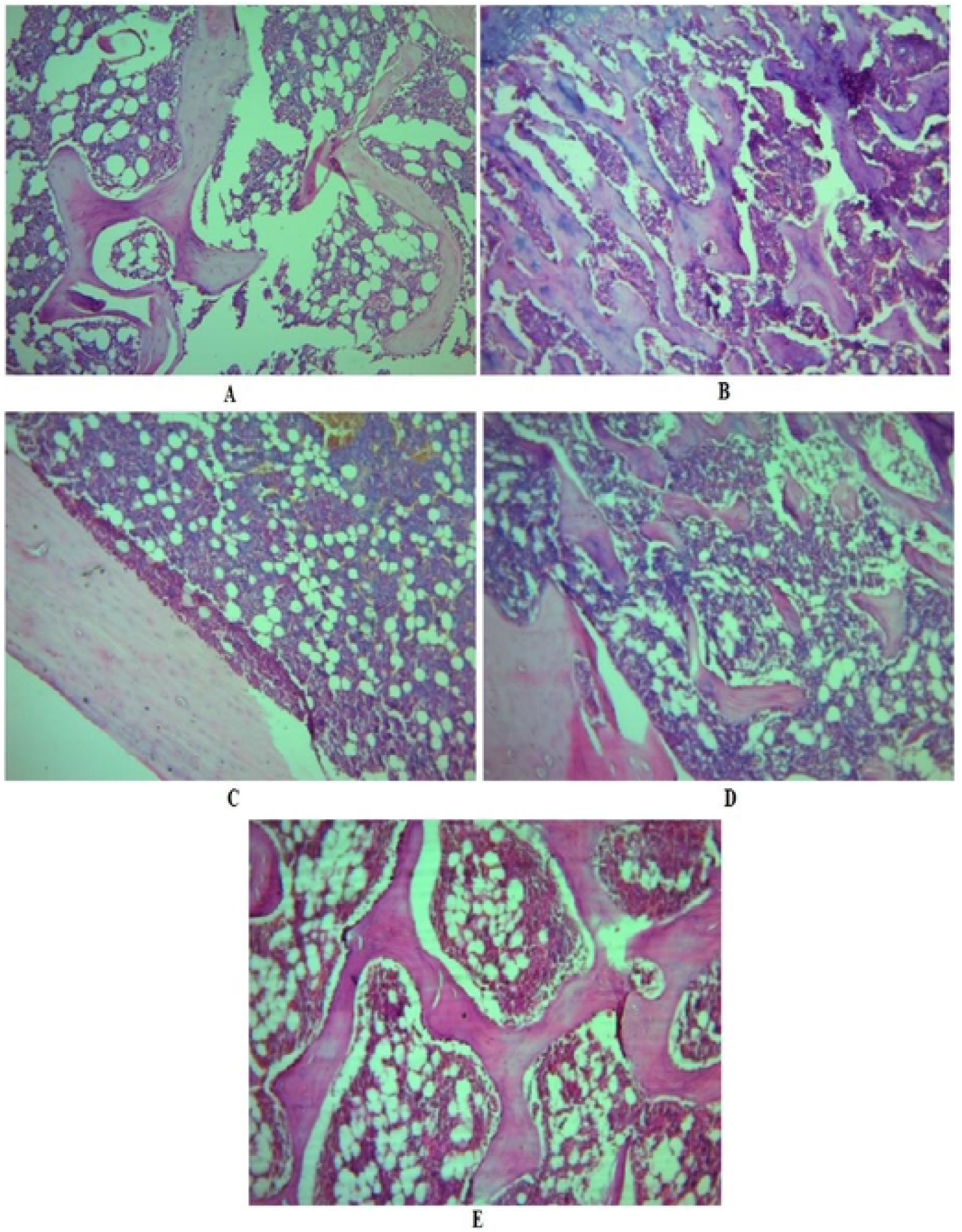
Effect of AEF of *P. tuberosa* on histopathology of the femur. **A,** Photomicrography of the femur of sham control group showing typical bone architecture; **B,** Photomicrography of the femur of OVX control group showing disruption of trabeculae; **C,** Photomicrography of the femur of raloxifene treated group showing improved trabecular thickness, and compactness of cells indicating mineralization of bone; **D,** Photomicrography of the femur of AEF-100 mg/kg treated group showing the improved trabecular thickness and bone architecture; **E,** Photomicrography of the femur of AEF-200 mg/kg treated group showing the restoration of typical bone architecture and increase in width of trabeculae.

#### *In vitro* cytotoxicity of AEF

Postmenopausal osteoporosis, which typically causes weakness of bone as the process of bone-resorption exceeds bone-formation because of estrogen-deficient state [33]. Hormone replacement therapy (HRT) is a choice to manage the problems in postmenopausal women, but continuous administration of HRT has the danger of cancer (breast, ovary and endometrial) development [34]. Therefore, we assessed the *in vitro* anticancer activity of AEF (31.5 - 500 μg/mL) in MCF-7 and MDA-MB-231 breast and SKOV-3 ovarian cancer cell lines. AEF displayed anticancer activity against the three cancer cell lines in a dose dependent manner (Fig 7) confirming that AEF is safe and can take care of the menopausal complications. AEF demonstrated better activity against ovarian cancer cells compared to breast cancer cells.

**Fig 7.**
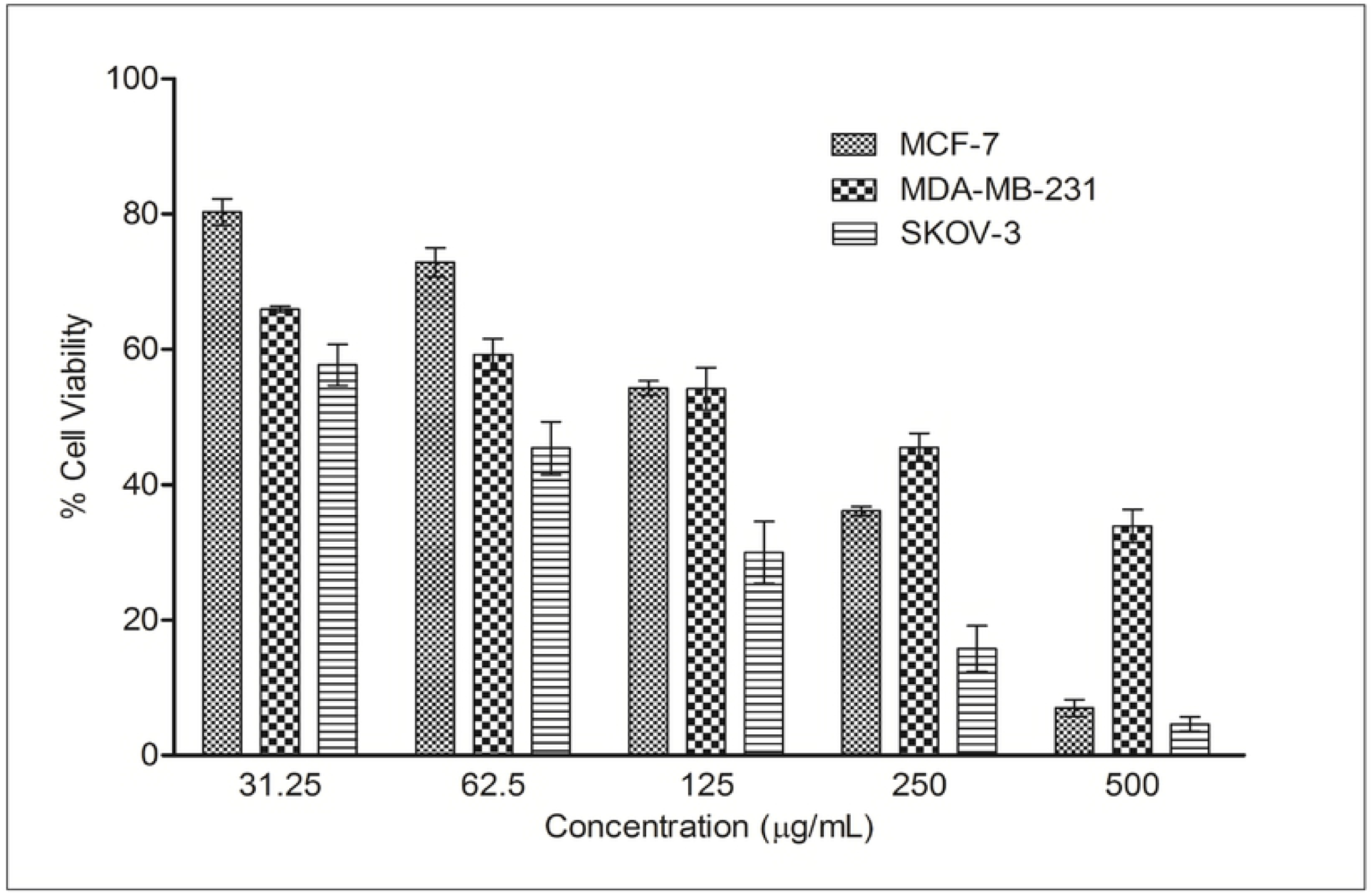
*In vitro* cytotoxicity of AEF of *P. tuberosa* against different cancer cell lines. Values are mean ± SEMs (n=3).

### Docking study

High performance thin layer chromatography analysis confirmed the presence of daidzein and genistein in AEF of *P. tuberosa* [31]. Docking pose of these phytoconstituents into estrogen receptor (ER) α (1 X 76) and β (1 X 7R) were evaluated and furnished in Fig 8 and 9. Genistein exhibited −26.1648 and −32.4084 docking score into ER-α and β active site, respectively. The docking score of daidzein into ER-α and β active site was −28.3129 and −31.8923, respectively.

**Fig 8.**
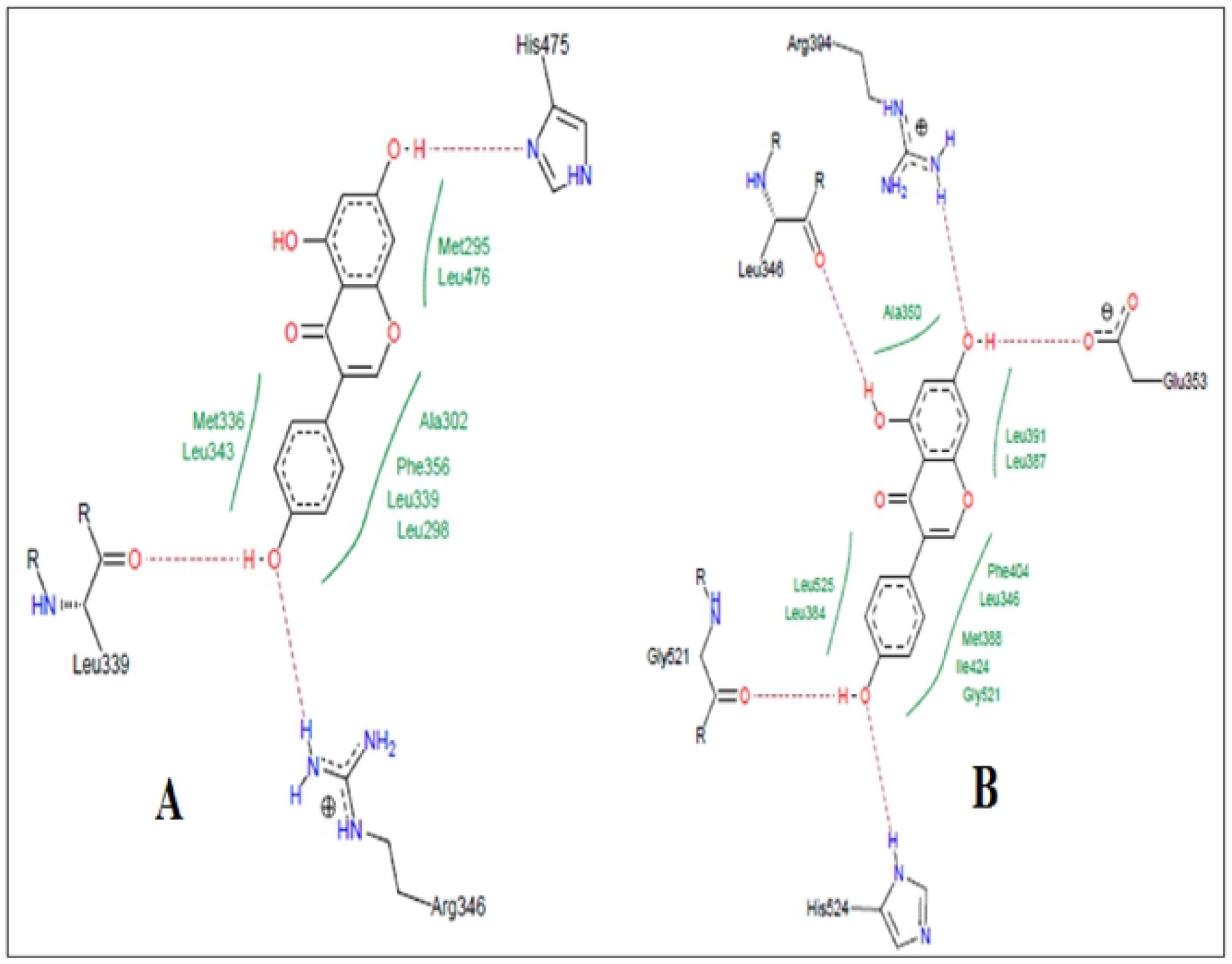
Docking study of genistein in estrogen receptors. **A,** Co-crystalized ligand of 1×76 (genistein) showing hydrogen bond with Arg346, Leu339, and His475. Hydrophobic interactions were also seen near the benzene rings with different amino acid residues of estrogen receptor α (PDB: 1×76). The ligand showed a docking score of −26.1648. **B,** Co-crystalized ligand of 1×7R (genistein) showing hydrogen bond with Leu346, Arg394, Gly521, Glu353 and His524. Hydrophobic interactions were also seen near the benzene rings with different amino acid residues of estrogen receptor β (PDB: 1×7R). The ligand showed a docking score of −32.4084.

**Fig 9.**
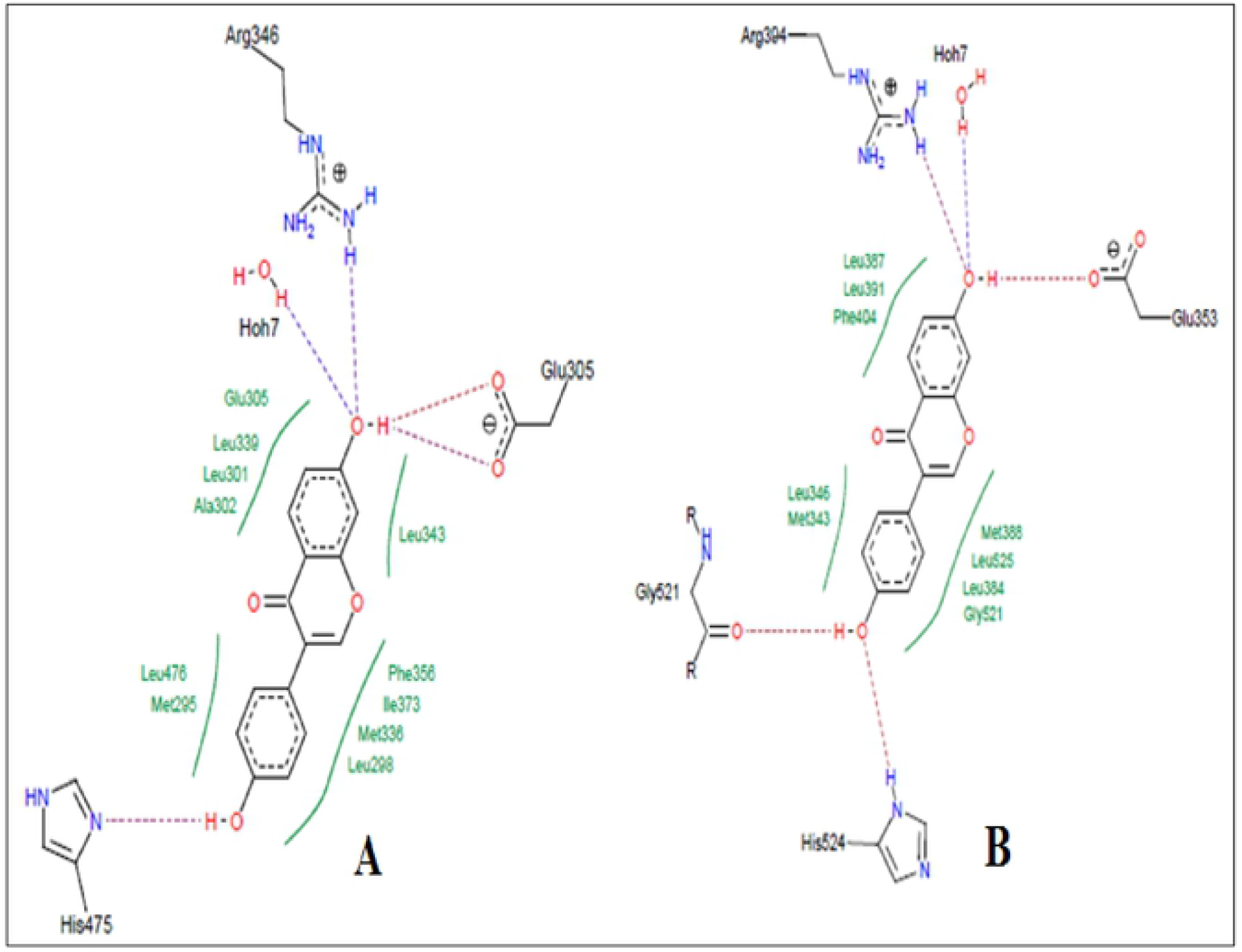
Docking study of daidzein in estrogen receptors. **A,** Docking pose of daidzein in estrogen receptor α (PDB: 1×76) active site with a docking score of −28.3129. Daidzein formed hydrogen bond with Arg346, Glu305, and His475. Hydrophobic interactions were also seen near the benzene rings with different amino acid residues of estrogen receptor α (PDB: 1×76). **B,** Docking pose of daidzein in estrogen receptor β (1 x 7R) active site with a docking score of −31.8923. Daidzein showed hydrogen bond with Arg394, Glu353, Gly521 and His524. Hydrophobic interactions were also seen near the benzene rings with different amino acid residues of estrogen receptor β (PDB: 1×7R).

## Discussion

Antioxidants play a major role in controlling the menopausal complications, including osteoporosis [7]. In this study, we have explored the *in vivo* anti-osteoporotic and *in vitro* anticancer activities of an AEF from the tubers of *P. tuberosa*. Ethanol extract of tubers of *P. tuberosa* and its various fractions (hexane, ethyl acetate, n-butanol, and aqueous) were analyzed for total phenolic and flavonoid content, and antioxidant activity. It was found that the ethyl acetate fraction contained maximum phenolic and flavonoid content, and antioxidant property, and was recognized the AEF.

The anti-osteoporotic activity of the AEF was evaluated in ovariectomized (OVX) rats by determining biochemical and biomechanical parameters, body and organ weights, and histopathology. The pattern of change in bone mineral parameters such as P and Ca in the present study confirms earlier findings of minor bone mineralization and balanced mineral homeostasis. The AEF did not change homeostasis and the effect might be because of enhanced absorption of calcium in intestine, as reported in previous studies [4, 7]. ALP and TRAP (bone turnover markers) activity are signs of bone osteoblast functioning and factors of bone formation. OVX increased these markers in serum because of the reduction in the estrogen level. HP is a commonly accepted biochemical parameter associated with bone metabolism, and its level is a sign of osteogenic activity. Urinary HP indicates break down of collagen due to high level of TRAP formed from activated osteoclast [35]. In the present study, the increased level of HP, TRAP, and ALP in OVX confirms reduced bone formation and an augmentation of collagen degradation. Further, administration of raloxifene and AEF reduced the above parameters, which indicates bone resorption inhibition property. AEF treatment produced positive effects on OVX-induced hyperlipidemia which could be due to existence of daidzein, genistein and β-sitosterol in *P. tuberosa*. Flavonoids could scavenge reactive oxygen species, which block TG secretion into the plasma and upset cholesterol catabolism into bile acids. Daidzein and genistein have been scientifically screened as antihyperlipidemic agents [36]. Further, presence of β-sitosterol in AEF hinders absorption of cholesterol by controlling lipogenesis and lipolysis [37].

Ovariectomized animal model, the most commonly used screening method of antiosteoporotic agents, has shown bone mineral density reduction leading to bone loss and increased susceptibility of fracture [38]. Healthy bones are normally compact and can tolerate considerable load. The compactness of the bone could be assessed by determining the bone strength. In the current study, the breaking strength of femur and 4^th^ lumbar vertebrae increased substantially by AEF of *P. tuberosa* proving the defensive effect of AEF against menopausal osteoporosis which are comparable to earlier reports [4, 7]. The phytoestrogens of AEF might have an estrogen like activity that manages osteoclast activity and reduces bone turnover.

The reduction of estrogen level in OVX animals causes increase in energy intake and elevated body weight [39]. Further, decrease in estrogen level due to OVX led to deposition of fat (as shown in the rise of total cholesterol and triglyceride) and hence an increase in body weight [11]. The observations in this study confirm that AEF administration reduced the level of cholesterol and TG in serum, which signifies the protective role of AEF against OVX-induced body weight gain. These observations corroborate the protective effect of AEF on adipose tissue and protection against the growth of osteoporosis [40].

Bone weakness is associated with bone mass, as well as its structure. Hence, histopathological analysis is a significant parameter to analyze the bone strength. OVX is associated with an increase in bone turnover, reduction in bone balance and loss in bone mineral density in the trabecular region of the femur [41]. The observed osteoprotective property of AEF manifested by superior trabecular architecture may be attributed to the secondary metabolites of AEF, which probably act as phytoestrogens to minimize bone loss [42].

In our earlier study, we reported the presence of two isoflavones, genistein and daidzein in the AEF [31]. In the present study, the phytoestrogenic nature of these two isoflavones was established by docking studies with estrogen receptor α and β, where both the compounds were found to have good affinity with both the receptors. As per earlier literature, estradiol has docking score of −18 and −17 into estrogen receptor α and β active site, respectively [40]. Our findings showed that bioactive compounds present in *P. tuberosa* have higher affinity compared to estradiol, which is also supported by earlier studies as daidzein and genistein showed high affinity into estrogen receptors [43]. Therefore, these two compounds might be mainly responsible for the antiosteoporotic property, which has been reported earlier [44, 45].

Phytoestrogens have been used as an alternative therapy for the management of menopausal osteoporosis as the regular use of hormone replacement therapy causes severe side effects, including cancer of breast and ovary [34]. Phytoestrogenic compounds also induce cell proliferation in ER-positive human breast cancer cells (MCF-7) [46]. However, in this study, AEF exhibited *in vitro* anticancer property in breast and ovarian cancer cell lines, suggesting its safety in the treatment of postmenopausal osteoporosis.

## Conclusion

The AEF from *P. tuberosa* contains bioactive compounds like genistein, daidzein, β-sitosterol, stigmasterol, etc. AEF exhibited marked antiosteoporotic activity in ovariectomy-induced osteoporosis. The protective effect of AEF might be attributed to its antioxidant potential as bone loss in osteoporosis could be due to generation of reactive oxygen species/oxidative stress along with other factors. Also, the presence of phytoestrogenic compounds such as daidzein and genistein in the AEF and their direct interaction with estrogen receptors may add to the protective effect. The AEF also exhibited significant anticancer activity in breast and ovarian cancer cell lines. The findings elucidated that AEF could be used as a safe therapeutics for controlling menopausal problems. However, further research is necessary for isolation of bioactive molecules from AEF and mechanistic studies are required to probe their antiosteoporotic effect.

## Acknowledgements

This work was supported by University Grants Commission, New Delhi, India [F.NO.5-63/2016(IC)]. The support of Dr. Pankaj Samuel, GC/MS Laboratory, Panjab University, Chandigarh, India, is highly appreciated for performing GC/MS analysis. We would like to express our sincere gratitude to Dr. Manik Ghosh, Department of Pharmaceutical Sciences and Technology, Birla Institute of Technology, Ranchi, India, for conducting docking studies.

## Supporting information

Results of the GC/MS analysis of the antioxidant enriched fraction (Table 3) are available as supporting information.

